# Random innervation of cerebellar Purkinje cells as a substrate for diverse representational learning

**DOI:** 10.64898/2026.06.26.734775

**Authors:** Adrian Holtrup, Ramin Khajeh, Wei-Chung Allen Lee

**Affiliations:** Department of Neurobiology, Harvard Medical School, Boston, MA, USA; Kirby Neurobiology Center, Boston Children’s Hospital, Harvard Medical School, Boston, MA, USA

## Abstract

Although the cerebellar microcircuit is among the most well-characterized systems for studying neural computation, recent connectomic analyses highlight deviations from canonical models, raising fundamental questions about connectivity, function, and learning in the system. A key feature of the circuit is the convergence of a vast number of parallel fibers (PFs) onto Purkinje cells (PCs). Prevailing models of cerebellar computation assume all-to-all connectivity at this intersection, whereby a single PC “samples” from all PFs. However, experimental evidence suggests that each PC is innervated by only a subset of accessible PFs. This partial sampling creates differential connectivity that may reflect an organizational principle of cerebellar learning, but its implications are poorly understood.

Based on electron microscopy (EM) reconstructions, we show that PF innervation of PCs is largely consistent with a Bernoulli model where connections are randomly and independently distributed within anatomical constraints. In further support of a random model, we observe that connections of the ascending branches of granule cells are not predictive of connections of their PFs, nor is connectivity correlated across separate spatial encounters of a PF with the same PC. In a model of the cerebellar circuit, we then address the possibility that partial connectivity is a substrate of learning, i.e., a fixed, random mask that enforces diversity between PCs. We find that when considering “ensembles” of PCs, random partial connectivity can indeed outperform all-to-all connectivity. Our results provide a theoretical framework for understanding the role of partial connectivity between cerebellar PFs and PCs and may have implications for cerebellum-like systems and beyond.

## INTRODUCTION

The vertebrate cerebellum has a long history as one of the most attractive neural circuits for studying how complex behavior emerges from organized neural networks (Albus et al., 1971; Cajal, 1888; Eccles, 1967; Golgi, 1874; Itō, 1984; Marr, 1969; Palay & Chan-Palay, 1974; Wolpert et al., 1998). Its appeal stems from a distinctive divergence–convergence architecture, which is characteristic of various other circuits across different species (Bell et al., 2008; Cayco-Gajic & Silver, 2019; Sawtell, 2017). At the input stage, mossy fibers (MFs) – afferents originating from diverse brain areas – enter the cerebellum and synapse onto granule cells (GrCs), the most numerous neuronal type in the brain (Andersen et al., 1992; Azevedo et al., 2009; Caddy & Biscoe, 1979; Herculano-Houzel et al., 2006). The axons of GrCs ascend he molecular layer of the cerebellar cortex where they bifurcate into long transversal branches, called parallel fibers (PFs). On their course through the molecular layer, dense beams of PFs are sampled by large Purkinje cells (PCs), the sole output of the cerebellar cortex (**Fig. 1a**).

**Fig. 1:**
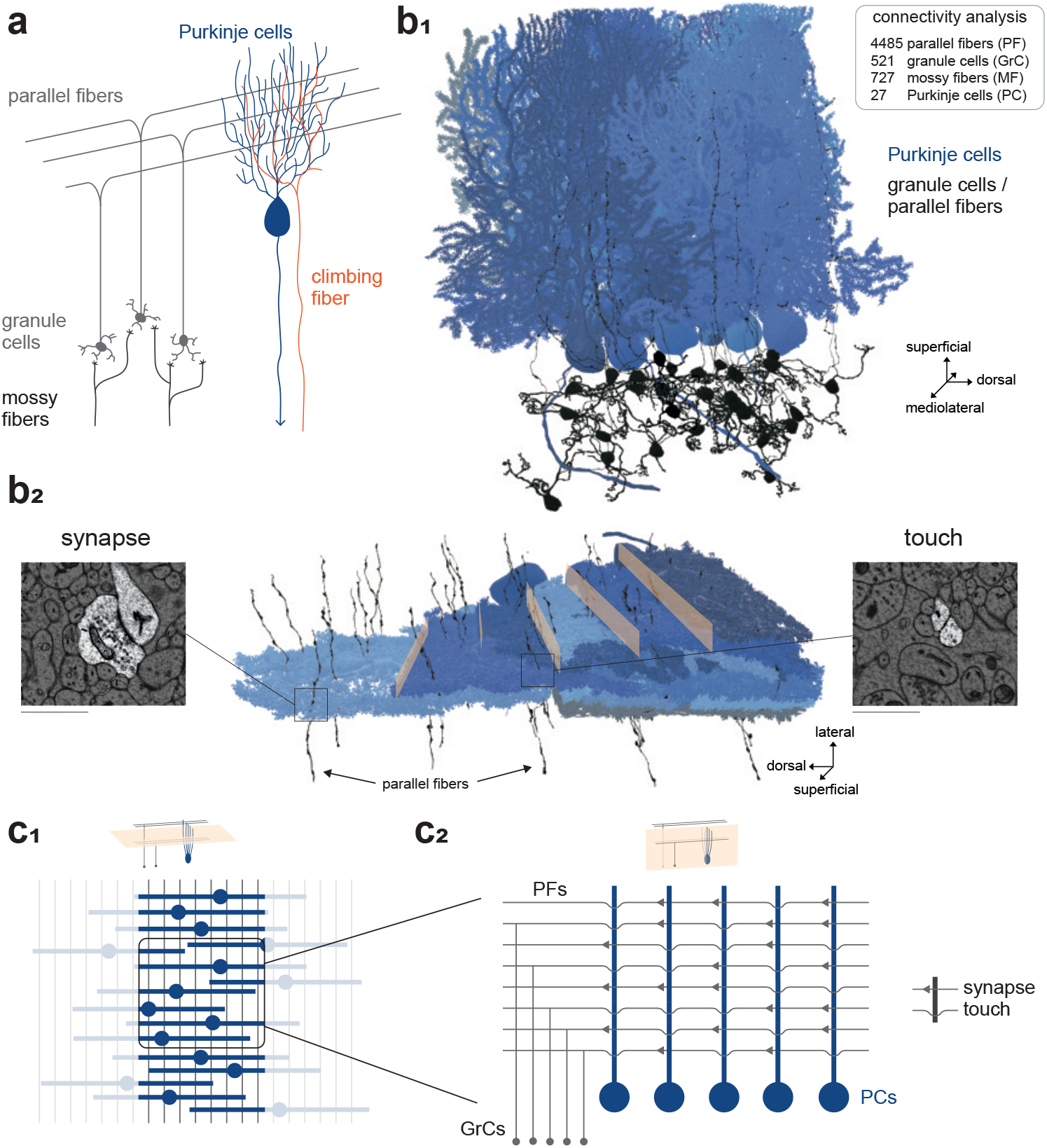
Convergence of parallel fibers onto Purkinje cells. **a)** Schematic of the cerebellar microcircuit. Granule cells (GrCs) receive inputs from mossy fibers (MFs) and send axons to the molecular layer, where they bifurcate into parallel fibers (PFs). PFs originating from local GrCs align with those from remote GrCs. In the molecular layer, Purkinje cells (PCs, blue) sample from PFs. Each PC is innervated by a single climbing fiber (orange). **b**_**1**_) EM reconstructions of a subset of GrCs and PFs, and PCs. The inset lists the numbers of analyzed cells. **b**_**2**_) EM reconstruction of PCs, illustrating the dense stacking of their sagittally-oriented dendrites. Dendritic trees are cut for better visualization of the layered architecture (indicated by the orange planes). Note how PFs penetrate the layers in the transversal direction. Zoomed-in EM images show PFs that either form a synapse onto a postsynaptic PC or merely “touch” the PC without a synaptic connection. Scale bar: 1 μm. **c**_**1**_) Schematic representation of PF sampling in the molecular layer. Dendritic trees of PCs span a plane orthogonal to the PF stream (gray). Individual PCs may sample diverse sets of PFs due to differences in spatial arrangement. However, we focus on differential connectivity among PCs that have access to the same set of PFs at the intersection of their dendrites. A cutout of such an intersection is highlighted. **c**_**2**_) Schematic of the wiring diagram from the cutout on the left. PFs touch all postsynaptic neurons but form synapses with (namely, connect to) a subset of them. Note that a connection always implies a touch. The empirical connection-to-touch ratio is called connection probability.

In cerebellar learning, an anti-Hebbian form of plasticity at PF-to-PC synapses shapes responses to particular PF activity patterns (Ito et al., 1982). Hence, connections from PFs to PCs are central to cerebellar learning. Interestingly, a considerable fraction of PFs pass within reach of PC dendrites without forming a synapse (Napper & Harvey, 1988; Nguyen et al., 2023; Park et al., 2023). This ‘partial connectivity’ gives rise to a non-trivial innervational pattern of adjacent Purkinje cells, unaccounted for by canonical theories of cerebellar learning (Albus et al., 1971; Litwin-Kumar et al., 2017; Marr, 1969). A thorough understanding of this partial wiring is still lacking.

To address this gap, we leverage electron microscopy (EM) reconstructions that map the cerebellar synaptic wiring diagram at the level of individual neurons (**Fig. 1a, b**) (Nguyen et al., 2023). We find that PF-PC connections are well described by a model in which each PC randomly and independently samples PFs within reach. This suggests that no select groups of PFs are systematically over- or under-sampled, nor are there groups of PCs characterized by specific PF innervation patterns. We next show that connections of GrC ascending branches are not predictive of the wiring of their parallel fibers, further supporting an unstructured model of GrC-PC connectivity.

We then extend canonical models of the cerebellar cortex to study the impact of partial connectivity on cerebellar learning. Previous work has well established that PCs are organized into sagittal zones, in which several PCs receive highly correlated climbing fiber inputs (Fakharian et al., 2025; Kostadinov et al., 2019; Llinas et al., 1974; Oscarsson, 1979; Zeeuw et al., 1997) and converge onto neurons in the cerebellar nuclei (Apps & Garwicz, 2005; Person & Raman, 2012; Sugihara et al., 2009; Zeeuw et al., 1994). This has long inspired the idea that groups of PCs, rather than individual PCs, serve as the functional unit of the cerebellum (Oscarsson, 1979). Combining both partial connectivity and Purkinje cell groups in a unifying framework, we propose that PF subsampling diversifies the inputs to otherwise redundant “ensembles” of PCs. In this modeling framework, we demonstrate that such partial PF-PC connectivity can be exploited to increase robustness in cerebellar learning tasks.

## RESULTS

To study cerebellar connectivity, we used EM reconstructions of GrCs and PCs from a previously acquired volumetric EM dataset of mouse cerebellar lobule V (Nguyen et al., 2023). PFs emanating from GrCs travel long distances in the molecular layer (Brand et al., 1976). Therefore, the dataset contains both local GrCs with their somas in the EM volume and remote PFs that originate from GrCs with somas outside the volume (Nguyen et al., 2023). Remote PFs are abundant, run perpendicular to PC dendritic trees (**Fig. 1b, c**), and typically form only one synapse per postsynaptic target (**Fig. 2e**). These properties make them ideal for investigating their wiring structure. Hence, we focused our analysis on remote PFs and compared our findings to local GrCs when applicable. For simplicity, we refer to remote PFs as PFs and reserve the acronym GrC for local GrCs including their ascending branches and parallel fibers.

**Fig. 2:**
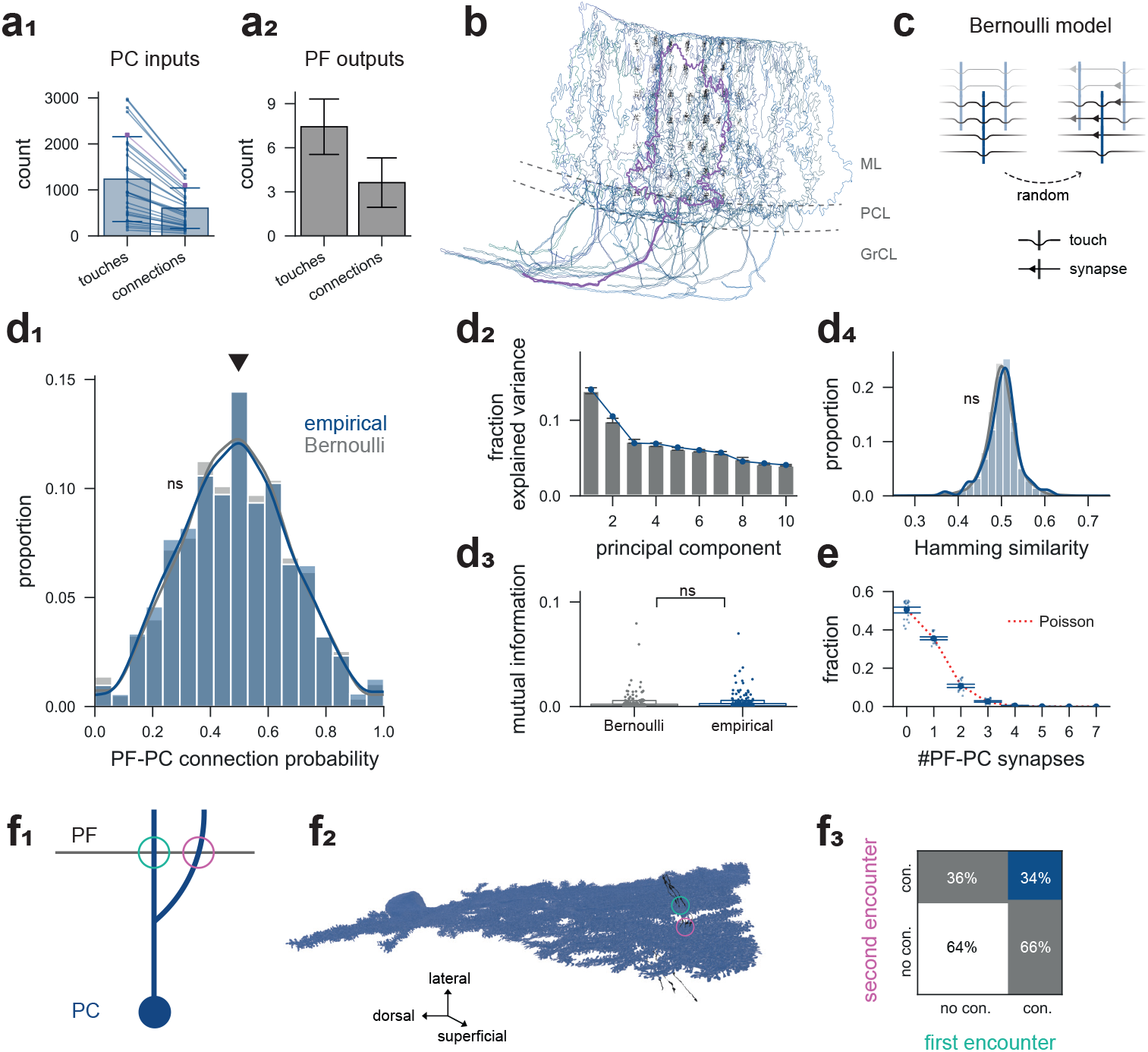
Anatomically constrained PF-PC connectivity is captured by a random Bernoulli model. **a**_**1**_) Numbers of touches and connections for the 27 PCs analyzed. Magenta squares correspond to the PC highlighted in **b** (error bars: mean ± SD). **a**_**2**_) Numbers of touches and connections for PFs analyzed (error bars: mean ± SD). **b)** Spatial outlines of the PCs analyzed, computed as α-shapes (Edelsbrunner et al. 1983), with an example PC highlighted (magenta, see **a**_**1**_). Black dots represent PFs, reconstructed following a grid pattern. ML: molecular layer; PCL: Purkinje cell layer; GrCL: granule cell layer. **c)** Schematic of the Bernoulli model. PF-PC touches are independently and identically converted into synapses, each with a probability equal to the empirical connection probability. **d**_**1**_) Distribution of empirical PF-PC connection probabilities compared with the Bernoulli model. The empirical distribution does not differ significantly from the Bernoulli model, indicating that no PFs are systematically over- or under-sampled by PCs (Monte Carlo test of variance, *p* = 0.12). Included are PFs with ≥4 touches. The distribution is centered around a connection probability of ∼0.5 (arrowhead). **d**_**2**_) Variance explained by the first 10 principal components of the PF-PC connectivity matrix for the empirical data (blue) and the Bernoulli model (gray). **d**_**3**_) Mutual information between pairs of PCs, computed on touch-constrained PF inputs. Mutual information is not elevated above baseline fluctuations (Monte Carlo test of mean, *p* = 0.36). **d**_**4**_) Distribution of pairwise PC Hamming similarity (Methods). PCs do not agree or disagree in their PF sampling more often than expected under the Bernoulli model (Monte Carlo test of mean, *p* = 0.16). Pairs of PCs with ≥30 shared PF touches are included. **e)** Synapse count distribution of PF inputs to individual PCs. The distribution is well explained by a two-parameter model combining Poisson-distributed “encounters” with a Binomial distribution conditioned on these “encounters” (red) (error bars: mean ± 95% CI). **f**_**1**_) Schematic of cases where a PF encounters dendritic branches of the same PC at two separate locations, allowing for a cross-validation analysis. **f**_**2**_) Example reconstruction of a PC (blue) and several PFs (black), corresponding to **f**_**1**_. **f**_**3**_) Mosaic plot of connectivity at the two separate encounters (Methods). Percentages indicate column proportions (e.g., the blue tile indicates that 34% of pairs connected at the first encounter are also connected at the second; correlated wiring would manifest as sizable differences in connected percentages across the two columns). Connections at the two encounters are uncorrelated, indicating that PF-PC connectivity is not governed by a systematic rule (868 PF-PC pairs involving 829 and 22 unique PFs and PC, respectively; Pearson *R* = -0.02).

Given the limited spatial span of a PC’s dendritic tree, only a subset of PFs in our dataset is accessible to each PC. To investigate how PCs sample from this subset, we computed “touches” between PFs and PC dendrites, i.e., a binary metric indicating whether the shortest distance between a PF and a PC dendrite falls below a set threshold (**Fig. 1b, Supplementary Fig. 1**, Methods) (Nguyen et al., 2023). The touch metric allowed us to calculate the conditional probability that, given an axon and dendrite touch, the two neurons form a synapse (connection). We refer to this value as the (empirical) connection probability.

### Parallel fibers converge in a Bernoulli-like manner

To better understand whether innervation of PCs by PFs follows a particular logic whereby certain PFs target specific PCs, we analyzed the structure of PF-PC convergence. We included only those PCs that were touched by at least 100 PFs (total reconstructed PFs in the dataset: 4,485). The median number of touches was 947 per PC, and each PC formed synapses with roughly 50% of its touches (mean connection probability: 49.5% ± 4% SD; **Fig. 2a**). Using this probability, we defined a Bernoulli null model in which each touch was converted into a connection with a probability equal to the empirical population value (**Fig. 2c**, Methods).

The Bernoulli model closely matched the distribution of empirical connection probabilities, indicating that no subset of PFs is systematically over- or under-sampled by PCs (**Fig. 2d**_**1**_, Monte Carlo test, *p* = 0.12). Similarly, principal component analysis of the connectivity matrix revealed no leading components that exceeded those of the Bernoulli model (**Fig. 2d**_**2**_) (Caron et al., 2013; Li et al., 2020). To assess correlations in sampling between PCs, we calculated mutual information (Shannon, 1948) on sampling of shared touches for pairs of PCs. Again, mutual information did not differ significantly from the Bernoulli model, suggesting that the sampling pattern of one PC provides little information about that of another (**Fig. 2d**_**3**_, Monte Carlo test, *p* = 0.36). Further, we found no elevation in Hamming similarity (**Fig. 2d**_**4**,_ Monte Carlo test, *p* = 0.16), increased shared input between pairs of PCs, or correlation of input vectors (**Supplementary Fig. 2b**). Minor deviations from the Bernoulli model include a small elevation in the variance of connection probabilities across PCs, suggesting some PCs may sample PFs slightly more densely than others (**Supplementary Fig. 2a**), and a slight decrease in PF-PC connection probability with increasing height in the molecular layer (Nguyen et al., 2023).

Extending the random model beyond binary connections, we found that the synapse count distribution of unitary PF-PC connections was well captured by a combination of a Poisson distribution – describing the number of encounters between a PF and a PC dendritic branch – and a Bernoulli model determining whether an encounter resulted in a synapse (**Fig. 2e**, Methods). This supports the hypothesis that synapses between a PF-PC pair form independently of other synapses between the same cells.

Independent synapse formation at different encounters of the same pair of cells would be a strong, orthogonal argument for non-selective wiring. To test this hypothesis, we isolated PF-PC pairs in which the PF came into close proximity with PC dendrites at two locations separated by at least 5 μm along the transversal axis (**Fig. 2f**_**1**_, **f**_**2**_, **Supplementary Fig. 3a**). Indeed, we found that the probability of a connection at the second encounter did not depend on the presence of a connection at the first (**Fig. 2f**_**3**_). This lack of correlation argues against selective wiring models in which connectivity between two cells is determined by cell identity.

**Fig. 3:**
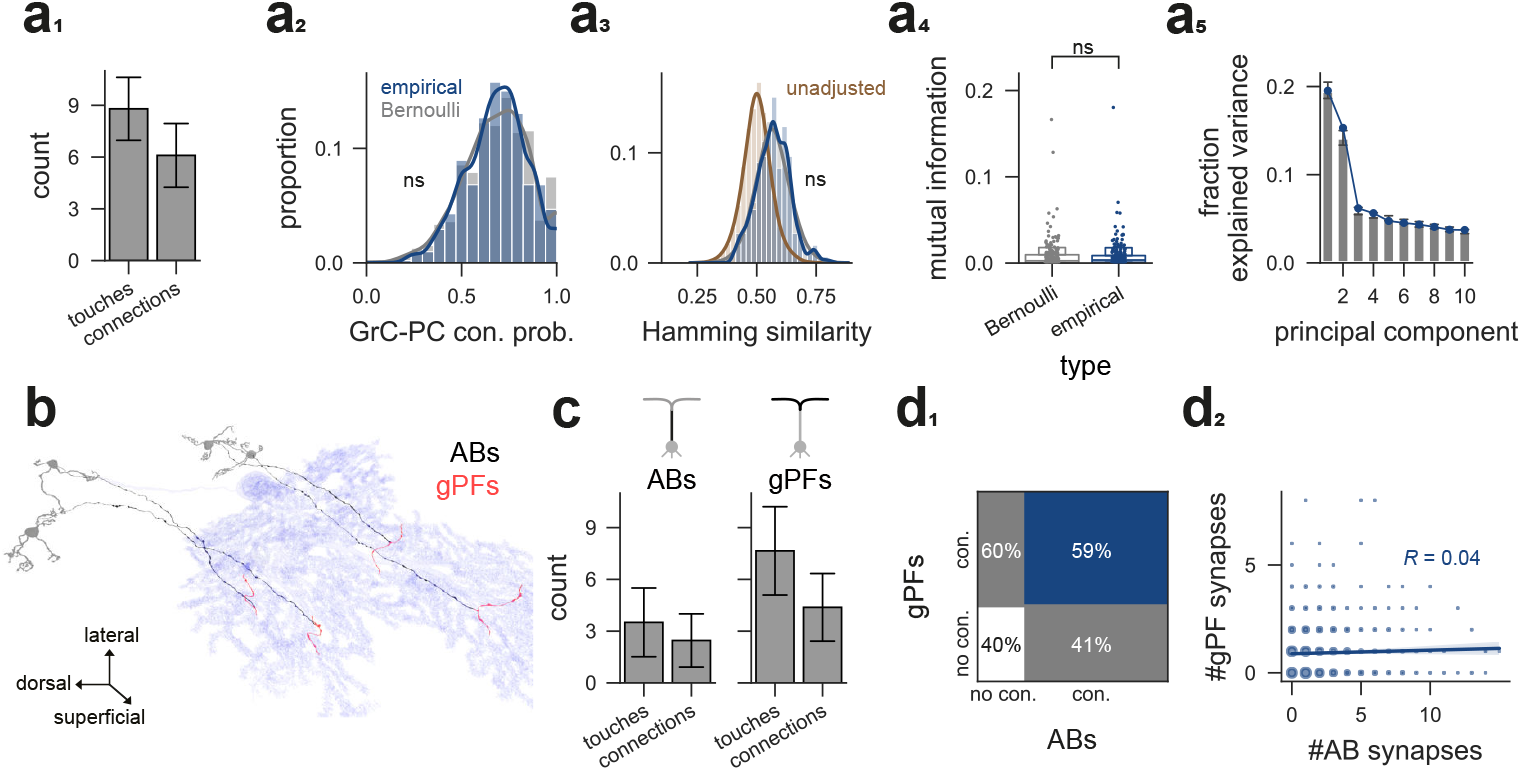
Ascending branches and parallel fibers of local granule cells wire independently. **a**_**1**_) Number of touches and connections for local GrCs (error bars: mean ± SD). **a**_**2**_) Distribution of empirical GrC-PC connection probabilities for local GrCs. The average connection probability is increased compared to remote PFs (0.69 vs. 0.5), yet the distribution agrees with a Bernoulli model adjusted to this elevated probability (Monte Carlo test of variance, *p* = 0.43). **a**_**3**_) Distribution of pairwise PC Hamming similarity for inputs from local GrCs. The Hamming similarity distribution is biased by the connection probability: when the corresponding Bernoulli model (gray) respects the elevated connection probability of local GrCs, it agrees well with the empirical distribution (blue; Monte Carlo test of mean, *p* = 0.23), whereas an unadjusted Bernoulli model with a connection probability of 0.5 shows smaller Hamming similarity (brown; Nguyen et al., 2023). **a**_**4**_) Mutual information for pairs of PCs with respect to their local GrC inputs (Monte Carlo test of mean, *p* = 0.59). **a**_**5**_) Variance explained by the first 10 principal components of the GrC-PC connectivity matrix for the empirical data (blue) and the Bernoulli model (gray). **b)** Example reconstructions of ascending branches (ABs, black) and parallel fibers of local GrCs (gPFs, red), touching a PC (blue). **c)** Analogous to **a**_**1**_ but separated for ABs and gPFs of local GrCs. **d**_**1**_) Mosaic plot of connectivity correlations between ABs and gPFs of local GrCs. Shown are cases where the AB and gPF both touch the same PC. Percentages indicate column proportions (e.g., the blue tile indicates that, of the cases where the AB connects to a PC touched also by the gPF, 59% of the time the gPF connects to the same PC, too). gPFs are not more likely to connect to the PC, provided the AB does so (Pearson *R* = -0.01). **d**_**2**_) Synapse counts of AB and gPF of a given local GrC onto PCs touched by both (Pearson *R* = 0.04).

### Ascending branches and parallel fibers of local granule cells wire independently

Local GrCs have been hypothesized to provide stronger and more focal excitatory drive to PCs than the numerous, but individually weaker unitary inputs from distant PFs (Bower & Woolston, 1983; Gundappa‐Sulur et al., 1999). We therefore next investigated the connectivity of local GrCs. The empirical GrC-PC connection probability is, on average, larger than that for PFs. This difference can be mostly attributed to their ascending branches (**Fig. 3a**_**1**_, **c**). Again, we created a Bernoulli model, now adjusted to the elevated connection probability. Like remote PFs, the connectivity of local GrCs largely agreed with the Bernoulli model (**Fig. 3a**). The adjusted model also reconciles a previously observed elevation in Hamming similarity for local GrCs (**Fig. 3a**_**3**_) (Nguyen et al., 2023).

A longstanding debate has concerned the different roles of the ascending branches of local GrC axons compared to their parallel fibers (Cohen & Yarom, 1998; Conti & Auger, 2024; Gundappa‐Sulur et al., 1999; Huang et al., 2006; Lu et al., 2009; Santamaria et al., 2002). To better distinguish how these different compartments contribute to GrC connectivity, we split local GrC axons into ascending branches (ABs) and parallel fibers (gPFs) (**Fig. 3b**). As expected from their ascending trajectory, ABs exhibited fewer touches but a relatively high connection probability (**Fig. 3c**). This is likely due to repeated encounters with the same PCs, as ABs also tend to form more synapses with each unique partner (**Supplementary Fig. 2f**_**2**_). In contrast, gPF connectivity was comparable to that of remote PFs (**Fig. 3c, Supplementary Fig. 2f**_**3**_).

The compartmentalization of GrCs into ABs and gPFs also allows for an analysis analogous to **Fig. 2f**: If wiring is cell-selective, gPFs should be more likely to connect to a given PC when the ascending branch of the same granule cell also connects to that PC. However, gPFs showed no such preference (**Fig. 3d**_**1**_). The synapse count of unitary AB-PC connections was not predictive of the synapse count of a gPF connection to the same PC either (**Fig. 3d**_**2**_), suggesting that connectivity of the two axonal compartments is independent. Collectively, these findings support a model of largely unstructured connectivity for both local GrCs and remote PFs.

### An ensemble model of Purkinje cells

Our analysis revealed that partial connectivity is a prominent feature of PF convergence onto PCs, but that this ‘subsampling’ is largely consistent with random, unstructured connectivity. The lack of distinct structure prompted us to investigate whether, and how, partial connectivity itself influences learning when integrated into canonical models of the cerebellum (Albus et al., 1971; Babadi & Sompolinsky, 2014; Cayco-Gajic et al., 2017; Cayco-Gajic & Silver, 2019; Kenyon et al., 1998; Litwin-Kumar et al., 2017; Marr, 1969; Xie et al., 2023). The model we consider consists of a mossy fiber layer of size *N* that encodes an input pattern *x*. This input is projected to a more expansive PF layer of size *M*, with *M* ≫ *N*, through the synaptic weight matrix *J*. The PF response is given by

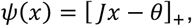

Where [·]+ is the rectified linear function and the threshold *θ* is chosen so that a fraction *f* of the PFs is active (see Methods). The vector *ψ(x)* is the PF-layer embedding of the input that PCs read out.

It is clear that with a single PC, partial connectivity is equivalent to shrinking the PF layer, namely omitting PF input and limiting information, which in general cannot benefit network performance. The situation is different if we consider not one, but a population of PCs, in which case partial connectivity allows different PCs to access distinct subsets of PFs, thereby diversifying their inputs. This is consistent with a series of experimental observations: Groups of PCs are arranged in sagittal zones where (1) they share highly correlated climbing fiber input (Groenewegen & Voogd, 1977; Llinas et al., 1974; Moatamed, 1966; Oscarsson, 1979; Schild, 1970; Sugihara et al., 2001; Zeeuw et al., 1997), (2) display simple spike synchrony (Bell & Grimm, 1969; Bell & Kawasaki, 1972; Fakharian et al., 2025; Schwarz & Welsh, 2001; Wise et al., 2010), and (3) converge onto the same cells at the deep cerebellar nuclei (Discussion) (Apps & Garwicz, 2000, 2005; Person & Raman, 2012; Sugihara et al., 2009; Zeeuw et al., 1994). These features suggest that groups of PCs rather than individual cells serve as the functional unit of cerebellar cortex.

We implement this feature in the model by considering an “ensemble” of Purkinje cells that are each connected to a random subset of the PFs. Crucially, this random mask remains fixed throughout learning. The probability that any given PF-PC pair is connected is defined by the parameter *ρ* (**Fig. 4a**, Methods). At *ρ* = 1, the ensemble consists of multiple identical PCs (all-to-all connectivity); values of *ρ* < 1 correspond to partial sampling of the PF layer (partial connectivity). The output of the *i*-th PC is then given by

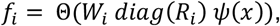

where *W*_*i*_ is the PF-PC weight vector, *R*_*i*_ ∈ {0, 1}^*M*^ is the random mask that determines the inputs to this PC, and Θ is the Heaviside step function. For now, the output of the ensemble, *f*_ens_, is defined as the majority vote over all PCs. Later, we investigate *f*_ens_ as the uniform average over linear PC outputs.

**Fig. 4:**
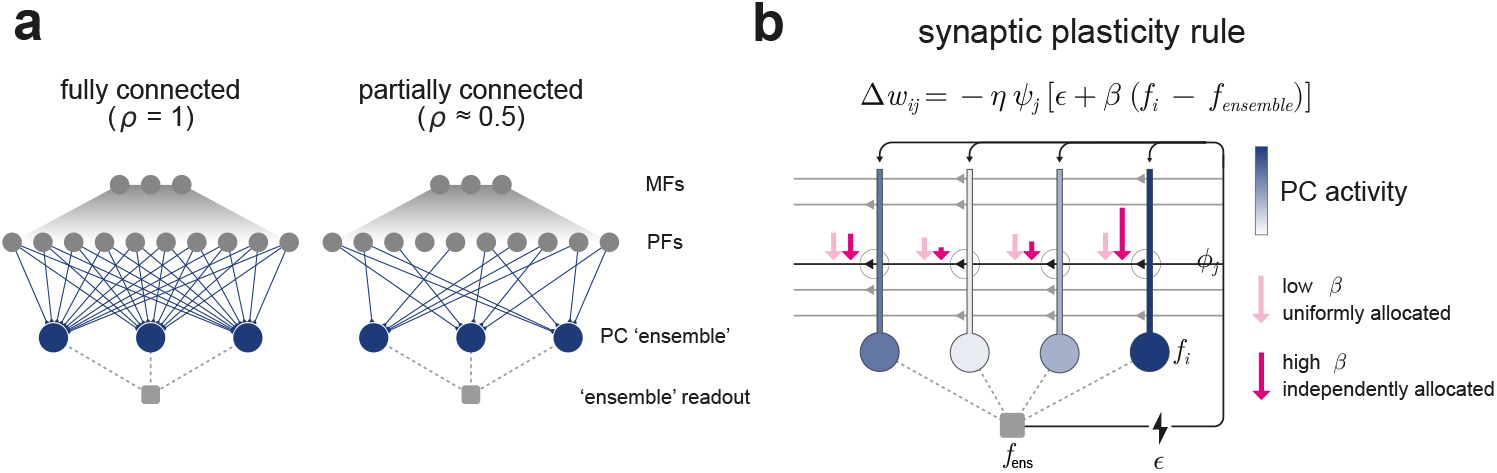
An ensemble model of Purkinje cells. **a)** Model architecture. Task inputs are embedded into the mossy fiber (MF) layer and projected to the expansion layer representing parallel fibers (PFs). An ensemble of Purkinje cells (PCs) receives input from PFs and produces an output by majority vote (classification) or by averaging (regression), which is then compared to a task-defined target. Each PC in the ensemble samples from a random subset of PFs; the size of this subset (i.e., the fraction of PFs connected) is set by *ρ*. Weights between PFs and PCs are subject to plasticity by an error signal. Importantly, only a single error signal is generated for the entire ensemble. **b)** Synaptic plasticity rule used for online learning. For ensemble learning, independence of the learners is critical. With a single error signal, this independence is achieved by differential susceptibility of ensemble members to that error. We implement this by introducing a third factor, *β*(*f*_*i*_ − *f*_*ens*_), which depends on individual PC activity (color indicates activity level). *β* controls the degree of differential susceptibility of PCs to the error, *ϵ*. Synapses onto the most active PCs undergo the strongest depression: larger magenta arrows indicate larger weight depression for the most active PCs.

In line with previous studies (Litwin-Kumar et al., 2017; Xie et al., 2023), we first evaluate our network on binary classification. PCs, in the canonical view, learn by instructive real-time feedback conveyed by climbing fibers, which leads to the depression of PF-PC synapses that were active shortly prior to climbing fiber input (Coesmans et al., 2004; Kostadinov & Häusser, 2022; Ohmae & Medina, 2015; Silva et al., 2024; Simpson et al., 1996; Zang & Schutter, 2019). This can be modeled as a conventional on-line anti-Hebbian plasticity rule, Δ*W*_*ij*_ ∝ − *ψ*_*j*_*(x)ϵ*, where *ϵ* represents climbing fiber feedback (error), with *ϵ* = 0 for correct outputs and *ϵ* = ±1 in the case of errors.

A hallmark of ensembles is that their members are trained individually; the benefit of the ensemble lies in its ability to mitigate errors made by a few members, provided that those errors are heterogeneous across the population (Krogh & Sollich, 1997). However, if PC responses are pooled to serve as a functional unit, a feedback signal must be generated from this pooled, joint output and is therefore shared by all ensemble members (see Discussion). As a result, the ensemble faces a challenge of maintaining diversity in light of a shared instructive signal.

To overcome this challenge, we introduce a third factor to the conventional anti-Hebbian cerebellar plasticity rule that depends on the activity of individual PCs (**Fig. 4b**):

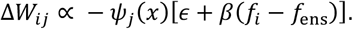

Since climbing fiber inputs evoke PF-PC LTD, we can assume that the ensemble output and the resulting climbing fiber signal are correlated during learning. In our learning rule, PCs with higher activity undergo stronger depression at their PF synapses, “punishing” them for their larger contribution to the feedback signal. This susceptibility to depression is controlled by the parameter *β*. At *β* = 0, the conventional rule is recovered, where PCs receive identical feedback signals. Larger values of *β* allocate the feedback signal in a way that depends on the individual PC activity, specifically on the deviation of individual PCs from the ensemble output (postsynaptic third factor). This is motivated by prior evidence showing that the extent (and sign) of PF-PC LTD is positively correlated with Ca^2+^ dendritic activity in PCs, which in turn is controlled by PF and inhibitory inputs (Bonnan et al., 2021; Eilers et al., 1997; Finch et al., 2012; Gaffield et al., 2018; Rowan et al., 2018).

### Diverse parallel fiber sampling can enhance learning

We began our investigation of network performance with a binary classification task, where the network was trained to learn random associations between mossy fiber inputs and binary labels and then tested on its ability to generalize to noisy versions of these patterns (**Supplementary Fig. 4a**) (Babadi & Sompolinsky, 2014; Cayco-Gajic et al., 2017; Litwin-Kumar et al., 2017; Xie et al., 2023). To isolate the diversity produced by subsampling, we first studied the setting in which PF-PC weights are initially uniform. In this case, ambiguity in the ensemble is due to differences in partial wiring alone. We found that, across different levels of generalization difficulty, partial connectivity outperformed full connectivity (**Fig. 5a**_**1**_). Analysis of performance as a function of *β* showed that the postsynaptic component in the learning rule is essential to exploit partial connectivity (**Fig. 5a**_**2**_). The advantage of subsampling can be understood as a tradeoff between the dimensionality of PF subsets and that of PC ensembles (**Fig. 5a**_**3**_) (Litwin-Kumar et al., 2017).

**Fig. 5:**
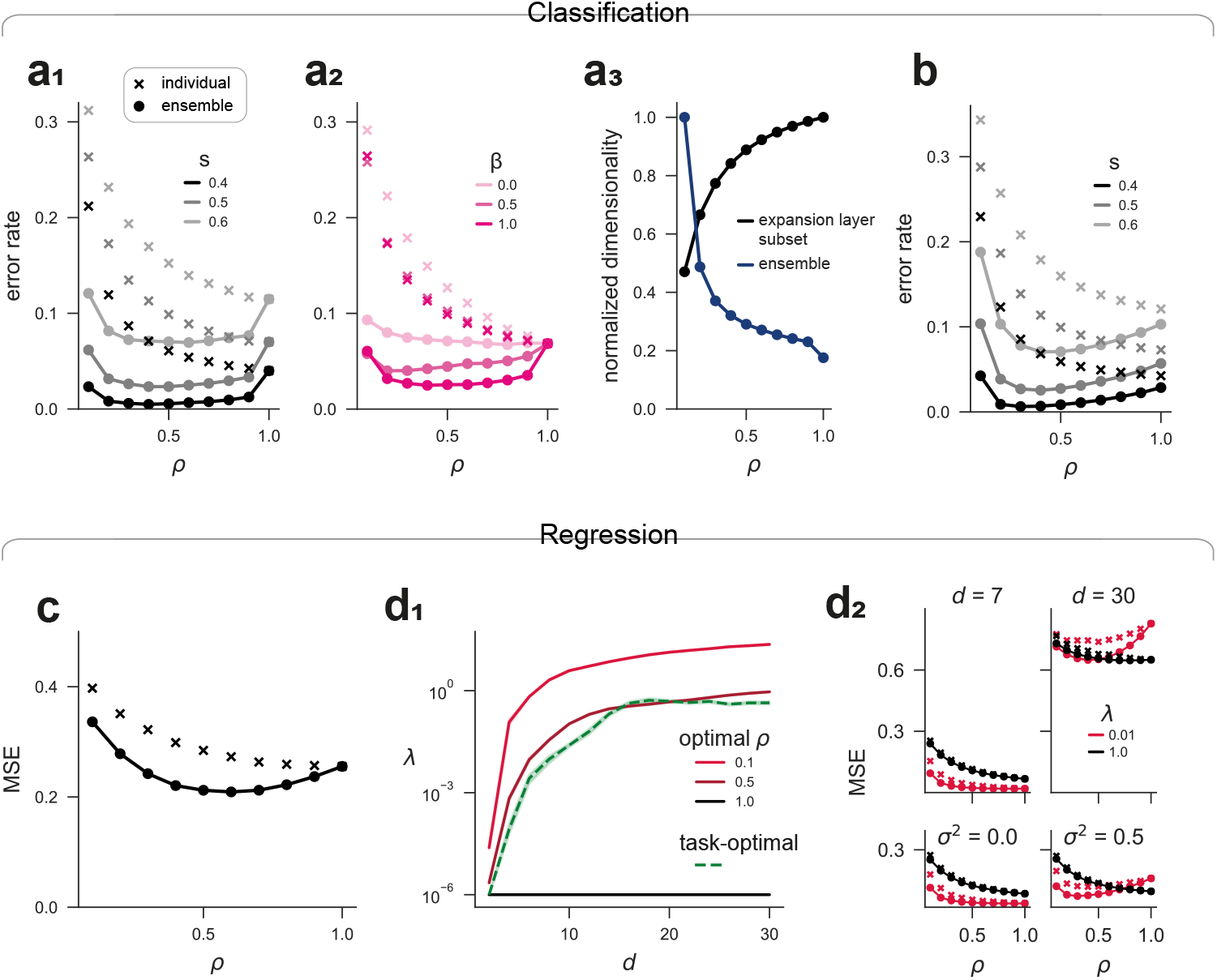
Diverse parallel fiber sampling can be exploited to enhance learning robustness. **a**_**1**_) Generalization error rate in a classification task with uniform weight initialization, as a function of *ρ*. Points: ensemble errors; crosses: errors of individual ensemble members. *s* controls the deviation between test and training patterns, i.e., the generalization difficulty. In this setting, any heterogeneity between the ensemble members derives from differential (partial) connectivity; for *ρ* = 1, ensemble performance is equal to individual performance. Note the increased performance for intermediate values of *ρ*. **a**_**2**_) Error rate as a function of *ρ* for different values of *β*, which sets the differential susceptibility of PCs to the error. This differential susceptibility is required to leverage the advantages of the ensemble. **a**_**3**_) Normalized average dimensionalities of PF subsets and PC ensembles as a function of *ρ*. Note that *ρ* mediates a tradeoff between ensemble and expansion-layer dimensionality. Normalization is done independently for ensemble and expansion-layer responses. **b)** Generalization error rate for ‘warm-start’ learning: Training starts with a weight matrix that has already stored unrelated patterns (after training on 50 prior, independent training sets). **c)** Example regression task using the on-line learning rule. Mean squared generalization error (MSE) as a function of *ρ*. Here, the fully connected network overfits the training data, resulting in suboptimal generalization performance. This is rescued by partial connectivity. **d**_**1**_) Adaptive implicit regularization of subsampling ensembles (ensembles with partial connectivity) in Ridge regression. Effective Ridge regularization penalty *λ* as a function of task dimension *d*. The green curve shows the task-optimal *λ*; black and red curves show the effective regularization implemented by networks with different *ρ*. Subsampling ensembles exhibit implicit regularization that adapts to the task: for *ρ* = 0.5, the induced regularization follows the task-optimal *λ* across a range of *d*, whereas the fully-connected network (*ρ* = 1) does not show this effect. Explicit Ridge penalty set to *λ* = 10^−4^. **d**_**2**_) MSE in Ridge regression as a function of *ρ*. Top: different task dimensions *d*. Bottom: different noise levels *σ* in the targets. *λ* denotes the explicit regularization penalty. With low explicit regularization (red), subsampling ensembles show more robust performance across task dimensions (top) and noise levels (bottom).

When then initialized with random weights, the ensembles still performed markedly better than the individuals, but the impact of partial connectivity was diminished and depended on the initial weight variance (**Supplementary Fig. 4c**). This was expected, as a random weight distribution provides an additional source of heterogeneity between ensemble members. We reasoned that weights that are initially diverse across ensemble members may rapidly converge when *ρ* is large, thereby reducing ensemble heterogeneity. To test this, we initialized the weights randomly and then trained the network repeatedly on different training sets without re-initialization, simulating an on-line learning setting more consistent with biology. In this case, partial connectivity preserved ensemble diversity and thereby improved performance, whereas with full connectivity, ensemble diversity was lost (**Fig. 5b**).

Finally, we evaluated the model in regression tasks, the second primary form of supervised learning (**Supplementary Fig. 4d**). While less established in a modeling context, regression tasks have recently been investigated in cerebellum-like architectures suggesting that they provide a more suitable framework for some cerebellar functions such as motor control (Xie et al., 2023). Building on this setup, we asked whether, and how, partial connectivity could be beneficial for performance. In this linear setup, we defined the PC response as *f*_*i*_ = *W*_*i*_ *diag(R*_*i*_*) ψ(x)* and set the ensemble output as the uniform average over ensemble members. The error was given by *ϵ* = *f*_*ens*_ − *y*, for a desired output *y* (see Methods).

In agreement with established results from machine learning studies (Atanasov et al., 2026; LeJeune et al., 2020), we found that partial connectivity introduces a task-dependent implicit weight regularization (**Fig. 5d**). This means that partial connectivity prevents the synaptic weights from adopting overly large values, thereby adaptively reducing the networks vulnerability to overfitting the training data. Assuming that optimally regularizing the full model is unfeasible under biological conditions, partial connectivity thereby increases the robustness of the network to changes in task configuration (**Fig. 5c, d**; **Supplementary Fig. 4e, f**; Methods).

## DISCUSSION

Although first observed decades ago (Napper & Harvey, 1988), the partial wiring between PFs and PCs has received little attention and remains largely enigmatic. We found that across different modalities of analysis, in the EM dataset, PF-PC connectivity appears largely unstructured when spatial distributions of the cells are accounted for. This holds true at the level of remote PFs as well as axons of local GrCs. In an ensemble model of cerebellar learning, we demonstrated that random, partial connectivity can lead to more robust model performance through diversification of individual PC responses.

### Random PF-PC connectivity

Key to the analysis of partial connectivity was the ability to characterize “touches” between PFs and PC dendrites. To define touches, one needs to impose a distance threshold between a presynaptic axon and a postsynaptic dendrite. This raises the question of what distance threshold is most appropriate to study the structure of PF-PC connectivity (Stepanyants et al., 2002). Choosing this threshold comes down to the question of what we learn about connectivity from knowing that a PF and a PC with a given minimal distance are not connected. Since we are interested in detecting potential cell-specific selectivity that may be rooted in physiological activity, the threshold should arguably lie within the range of PC spine growth. It is therefore reasonable to choose a touch-threshold below ∼1 μm. Here, we use a threshold of 160 nm for most analyses but examined the effects across different thresholds (**Supplementary Fig. 1**).

While many statistics are precisely reproduced by the Bernoulli model, some moderate but distinct deviations persist. Specifically, PF-PC connection probability depends weakly on height in the molecular layer (Nguyen et al., 2023), and PCs show a slightly elevated variance in connection probability. Both effects are linked to the morphology of the PC dendritic trees (**Supplementary Fig. 2a**). Given the many biological factors that can weakly influence connectivity (e.g., differences in the angulation of PC dendritic trees) and the limitations of our touch metric (e.g., touches onto thick dendritic branches where PFs are unlikely to form synapses), it is not surprising that some measures are not fully captured by the Bernoulli model. Thus, it seems unlikely that these deviations reflect deeper underlying structure.

A recent study has proposed that, during development, pairs of PCs share less PF input than predicted by chance (Dhanyasi et al., 2025), an observation that does not match our findings in the adult mouse. This may reflect ongoing developmental processes. However, this finding is noteworthy from a combinatorial perspective: PCs sampling from a shared pool of PFs have very limited leeway to undershare inputs in a pairwise manner, unless the number of PCs or their connection probability is small. In Dhanyasi et al. (2025), the distance to determine potential synaptic partners (analogous to touches) is on the order of microns. If too many cells are included in the space of potential connections, shared input may be underestimated and the variance of shared overlap for pairs of PCs is elevated (**Supplementary Fig. 1**). Future studies will be needed to determine the degree to which PC innervation patterns differ across developmental stage, cerebellar regions, individuals, and species.

### Functional implications

We can distinguish three interpretations of the results of our connectivity analysis:

1. Partial connectivity is indeed unstructured (i.e., the network is functionally invariant to reshuffling individual connections within constraints) and is a purposeful feature of the cerebellar cortex. This suggests the role of partial connectivity may be a substrate, as opposed to a result of learning. This is the interpretation that we focused on here and one that naturally motivates an ensemble framework, although it is not a necessity for our ensemble model.
2. Partial connectivity is indeed unstructured, merely reflects anatomical constraints and offers no functional benefit. This implies that a natural bound on the total number of inputs for each PC exists. This possibility cannot be excluded, but would require reconsidering the effective PF expansion ratio, or otherwise describe how clusters of PCs cooperate. In general, our arguments about diversification of PCs through partial connectivity would still apply under this scenario.
3. Partial connectivity is structured, (i.e., the network is not functionally invariant to reshuffling individual connections), but the underlying structure is such that it goes undetected with our analysis. Since it is impossible to prove that connectivity is unstructured, we cannot exclude this possibility. However, we can ask if there is a plausible and parsimonious hypothesis that aligns empirical connectivity with structured wiring:

While there are not many functional interpretations of partial PF-PC connectivity, previously, it has been suggested to be the result of learning-induced synaptic pruning (Aziz et al., 2014; Morizawa et al., 2022; Wang et al., 2014 but see Black et al., 1990; Ho et al., 2021). In this view, what we observed as touches would correspond to eliminated synapses and the connectivity would be structured by the definition given above.

The theoretical motivation for this idea stems from the work by Brunel et al. (2004) showing that at maximum storage capacity at least half of PF-PC synapses are silent (**Supplementary Fig. 5c**). It is important to distinguish between “touches” and the notion of “silent synapses”. In our analysis, a touch is purely a spatial relationship. By contrast, silent synapses are synapses that are electrophysiologically undetectable or non-functional (Isope & Barbour, 2002). Thus, our findings do not contradict predictions about the prevalence of silent synapses. However, if initially silent synapses were subsequently eliminated, touches would in fact be an expression of former silent synapses.

We performed simulations to assess whether the observed connectivity could be reproduced by pruning silent synapses following information storage. In this model, PCs initially share PF input according to their empirical touches (**Supplementary Fig. 5**). We found that correlations in climbing fiber input introduce similarity in PF sampling between PCs. Depending on the geometry of the PF code, even without shared climbing fiber input, such correlations in the modeled connectivity persist (**Supplementary Fig. 5d, e**). Together with the evidence that the fraction of silent synapses is likely much larger than 50% (Ho et al., 2021; Isope & Barbour, 2002) and the requirement that pruned synapses must remain as “potential” connections, it seems unlikely that partial PF-PC connectivity is the expression of a stored memory.

Further, if partial connectivity is a result of learning and, hence, dependent on PF activity and cimbing fiber input, one would expect correlated connectivity at repeated encounters of a given GrC/PF with dendrites of the same PC. Such a correlation was absent in our analysis (**Fig. 2f, 3d**). If connectivity was heavily determined by a local postsynaptic signal, correlations between different dendritic branches of the same PC could be less pronounced. Yet, given that such branches are typically innervated by the same climbing fiber, an entire lack of such correlation would still be unexpected.

Although this is only one example scenario in which connectivity derives from learning, our findings point to a more general discrepancy between the correlations induced by shared innervation of PCs and the lack of structure in the empirical connectivity. Hence, any theory in which partial connectivity is learning-induced must explain how correlations are hardly imprinted in the network despite shared PF and correlated climbing fiber input of adjacent PCs.

### Ensemble model

Established theories of cerebellar learning do not account for the fact that PCs sample only a fraction of PFs within reach. The lack of discernible structure in PF-PC wiring motivated us to explore subsampling as a substrate for, rather than a product of, learning. Since *a priori* subsampling for individual PCs can only lead to information loss, this naturally prompted us to consider ensembles.

An ensemble framework is reasonable only if there exist groups of PCs that subserve the same task. Of particular importance for our ensemble framework are functional experiments showing that climbing-fiber-evoked responses in PCs are highly co-tuned within sagittal zones (Ikezoe et al., 2023; Kostadinov et al., 2019) and that this co-tuning provides a principled basis for grouping PCs (Fakharian et al., 2025; Herzfeld et al., 2015). The anatomical substrates of this co-tuning are likely electrical coupling and shared input between the inferior olivary neurons as well as shared climbing fiber innervation between PCs within sagittal zones (Groenewegen & Voogd, 1977; Llinas et al., 1974; Moatamed, 1966; Oscarsson, 1979; Schild, 1970; Sugihara et al., 2001; Zeeuw et al., 1997). In further support of an ensemble framework, adjacent PCs display simple spike synchrony (Bell & Grimm, 1969; Bell & Kawasaki, 1972; Fakharian et al., 2025; Schwarz & Welsh, 2001; Wise et al., 2010), and converge at the level of deep cerebellar nuclear neurons with an estimated ratio of 34:1 – 52:1 (Apps & Garwicz, 2000, 2005; Person & Raman, 2012; Sugihara et al., 2009; Zeeuw et al., 1994).

In machine learning, it is well established that averaging over diverse subnetworks helps capture meaningful patterns in the data while ignoring idiosyncratic ones, a fundamental tradeoff when learning from examples (Atanasov et al., 2026; Breiman, 1996; Bryll et al., 2003; Geman et al., 1992; Hinton et al., 2012; Krogh & Sollich, 1997; LeJeune et al., 2020; Patil & Du, 2023; Patil & LeJeune, 2024; Srivastava et al., 2014; Wager et al., 2013; Yao et al., 2021). In line with this, our computational analyses show that subsampling PFs creates diversity in the PC layer that helps mitigate errors of individual predictors and can outperform fully connected networks.

Crucially, as we have shown, the ensemble mechanism requires decorrelating the training of individual PCs. In a real-time setting, however, feedback arising through interactions with the environment, for instance, cannot be easily assigned to each individual PC, making such decorrelation non-trivial. We suggested a solution to this problem based on differential susceptibility of PCs to a common teaching signal. This was achieved by incorporating the activity of PCs (postsynaptic) into the learning rule of the model.

The postsynaptic component of this learning rule is supported by studies in cerebellar slices (Bonnan et al., 2021; Eilers et al., 1997; Finch et al., 2012; Gaffield et al., 2018; Rowan et al., 2018). The learning mechanism also provides a natural explanation for the recent finding that inhibition of PCs by molecular layer interneurons reduces the susceptibility of PF-PC weights to climbing-fiber-mediated LTD (Rowan et al., 2018). Our learning rule allows predictions about how PF-PC synaptic plasticity is jointly governed by individual PC activity, population-level PC activity within sagittal zones, and climbing fiber input. Specifically, it predicts that LTD would be particularly pronounced at PF synapses on PCs with high simple spikes rates within their population.

Further, our plasticity rule depends on the deviation of the individual PC response *f*_*i*_ from the ensemble output *f*_ens_. The latter is a non-local term, raising the question of how it is made available to PF-PC synapses. If such a mechanism exists, a natural candidate for conveying this information is the climbing fiber signal. While the composition and computation of the climbing fiber signal are largely enigmatic, the multiple loops between PC output and the inferior olive make this a biologically feasible option (De Zeeuw & Ruigrok, 1994; Ruigrok & Voogd, 1990). We would then expect ensemble activity to influence the climbing fiber signal in two ways; through its impact on the error signal *ε* and through the contained term *βf*_ens_. Alternatively, the comparison between individual PCs and their ensemble partners could be mediated through direct collaterals in the cerebellar cortex (Lackey et al., 2026; Osorno et al., 2022; Witter et al., 2016). Yet it is also possible that variants of the suggested learning rule exist that, provided a suitable task, do not require the comparison between individual and joint output.

The behavior we observe in regression tasks agrees well with machine learning studies demonstrating analogies between subsampling and explicit weight regularization. This fundamental technique, used in training neural networks, prevents weights from becoming overly large and thus overfitting the training data (LeJeune et al., 2020; Patil & Du, 2023; Patil & LeJeune, 2024; Yao et al., 2021). In fact, the proper level of weight regularization depends on the data to be learned and is therefore generally unknown. This may be especially important in a biological context where diverse, noisy data must be processed in real time. Hence, wherever biological networks face the threat of overfitting, subsampling may be a natural and efficient alternative to regularize model complexity in an adaptive manner.

An interesting extension of our model would be to incorporate a learned, rather than uniform, pooling of the PCs (Krogh & Sollich, 1997). In this view, the weights of the PCs onto cerebellar nuclei neurons would undergo plasticity (Ohyama et al., 2006). Similarly, one could study further means of diversifying the PC ensembles. A concurrent study suggests diversifying PCs by coupled but heterogeneous climbing fiber input (Ruben & Pehlevan, 2026). This approach is related to subsampling in sample space, whereas our model uses feature subsampling.

Future work is needed to determine whether the innervation of remote PFs differs across microzones. The volume studied here has a sagittal extent of 49.5 µm and is therefore likely to lie within a single microzone (roughly 100-200 µm wide; (Oscarsson, 1979). Addressing PF connectivity across adjacent microzones, which receive distinctly tuned climbing fiber inputs, will require extending the ensemble framework presented here. Such efforts will help uncover a more comprehensive logic of parallel fiber innervation that supports cerebellar learning.

## METHODS

### Serial EM

All EM data along with reconstructions of MFs, GrCs, PFs, and PCs used in this work are from a previously imaged 776 μm X 753 μm X 49.5 μm volume of lobule V consisting of 1,167 sections of a P40 male mouse cerebellum (Nguyen et al., 2023). The synapse prediction (*synful*) had previously been applied to the whole dataset and evaluated on a sample basis (F-score: 0.938, Nguyen et al., 2023).

### Mesh reconstructions and touches

To control for spatial factors confounding the connectivity analysis, we calculated “touches” between GrCs/PFs and PCs. To that end, mesh representations with 16 nm resolution were generated for PCs, GrCs, and PFs. For local GrCs, we computed meshes for (1) the soma plus dendrites and (2) the axon and further subdivided the axon into ascending branch and parallel fiber. This allowed us to analyze touches for the different components of the GrC axons separately. Pairs of cells were defined as touching if their closest distance was no larger than 160 nm (Nguyen et al., 2023). Sensitivity analyses largely suggested the consistency of our findings across thresholds with certain expected, systematic deviations for large threshold values (**Supplementary Fig. 1**).

Outlines of PC dendritic trees in **Fig. 2b** were computed as α-shapes (Edelsbrunner et al., 1983) using the Python *alphashape* package.

To conduct a cross-validation analysis of PF-PC connectivity for individual pairs (**Fig. 2f, Supplementary Fig. 3**), we selected PF-PC pairs for which two separate locations existed where the PF was within 160 nm of a dendritic branch of the PC (touch-defining). The two locations were considered separate if a gap of at least 5 μm in the transversal direction existed between them, throughout which the distance between the PF and the PC exceeded 160 nm.

### Connectivity analysis

Unless stated otherwise, GrC/PF-PC connectivity was analyzed as binary. For connectivity analyses that involved GrC/PF-PC connection probabilities (**Fig. 2, 3**), unless mentioned otherwise, a threshold of ≥ 4 PC-touches was applied, below which GrCs/PFs were excluded.

Principal component analysis (**Fig. 2d**_**2**_, **Fig. 3a**_**5**_) was performed on the feedforward connectivity matrix where rows correspond to postsynaptic neurons (Caron et al., 2013; Li et al., 2020). Empirical pairwise mutual information (**Fig. 2d**_**3**_, **Fig. 3a**_**4**_) between PCs was calculated as 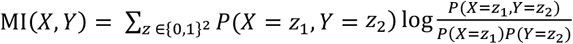 where *X* and *Y* are the binary connection vectors of the respective PCs onto their shared touches. Hamming similarity (**Fig. 2d**_**4**_, **Fig. 3a**_**3**_) for a pair of PCs was calculated as the fraction of agreement across all shared PF-touches, i.e., cases where both PCs either connect or both do not connect: 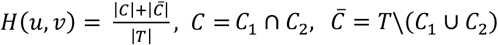, where *C* is the set of PCs that both PFs *u* and *v*, connect to, *T* is the set of their shared touches and 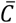 is the set of PFs within their shared touches that none of them connect so. Similarly, relative overlap (**Supplementary Fig. 1, 2c**) was defined as 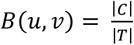.

### Random and structured connectivity models

Bernoulli connectivity models were built as PF-PC adjacency matrices, where each entry between a PF and its touches was drawn *i*.*i*.*d*. from *Bernoulli(p). p* here denotes the mean empirical connection probability of PF-PC touches. Structured scenarios were created based on information storage in the PCs and subsequent pruning of silent synapses (see *Network simulations*).

In **Fig. 2e** and **Supplementary Fig. 2d**, we show a random model that reconstructs the synapse count distribution for individual GrC/PF-PC touches. We define the probability that *k* synapses are formed between a touching PF-PC pair as 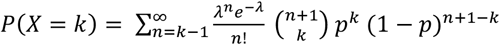, where *X* is the number of synapses, *λ* + 1 is the expected number of encounters between the PF and PC dendrites, and *p* is the Bernoulli probability that a synapse is formed at an encounter. Shifting *n* by 1 is a correction reflecting the fact that we consider only touching pairs. Otherwise, we would obtain a simple Poisson with parameter *pλ*. The optimal fit was determined numerically by least-squares-optimization.

In **Supplementary Fig. 2a**_**1**_, we consider a Bernoulli model that is adjusted for the slightly reduced PF-PC connection probability in the more superficial part of the molecular layer (Nguyen et al., 2023). In this model, the connection probability of a given PF is determined by its height in the molecular layer, based on a least-squares linear regression of empirical connection probabilities against height.

Another adjusted model is shown in **Supplementary Fig. 2c**_**1**_. Here, instead of using the mean population connection probability, we apply the individual probability of each PC to model the relative overlap.

### Network simulations

#### Network backbone

All networks consist of an input layer of size *N*, an expansion layer of size *M*, and, when applicable, a readout layer of size *Q*. Task inputs of dimension *d* are embedded into the input layer by a linear transformation *A* ∈ ℝ^*N*×*d*^, where *A* is a random matrix sampled from the Haar group (Muscinelli et al., 2023). Input and expansion layer are connected by a weight matrix *J*, with 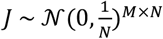. The expansion of input *x* is defined as *ψ(x)* = *ReLU(Jx* − *θ). θ* is chosen depending on *x* so that fraction *f* of *ψ(x)* is nonzero. Ensembles *W* ∈ ℝ^*Q*×*M*^ with Bernoulli connection probability *ρ* are created by linear masks *R* ∼ *Bernoulli(ρ)*^*Q*×*M*^. Let *W*_*i*_ ∈ ℝ^*M*^ be the *i*-th ensemble member. Its input is then *W*_*i*_ *diag(R*_*i*_*) ψ(x)*. In supervised learning tasks, we considered ensembles of size 20.

#### Information storage

In **Supplementary Fig. 5**, we compare the empirical connectivity to that of a model where patterns are stored in PCs and their synapses are pruned afterwards (Brunel et al., 2004). Initially, PCs were connected to all PFs that they touch. Hence, pairs of PCs shared PFs as determined by their touches. The learning protocol consisted of storing associations between PF activity patterns and binary labels. The similarity between PC labels, and thereby the similarity in the stored associations, was controlled by a parameter *s*. For *s* = 0, labels are identical for all PCs (shared climbing fiber); for *s* = 1, they are fully independent (different climbing fibers). Correlated labels reflect the fact that adjacent PCs receive highly similar climbing fiber input (Groenewegen & Voogd, 1977; Ikezoe et al., 2023; Kostadinov et al., 2019; Llinas et al., 1974; Moatamed, 1966; Oscarsson, 1979; Schild, 1970; Zeeuw et al., 1997). After learning, the 50% smallest weights were considered disconnected. We trained the models using a perceptron learning rule independently for each PC. PC weights were forced to be strictly nonnegative. The PC output was determined as 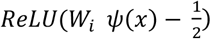.

Further, we built a model consisting of 20 artificial PCs with equal inputs but unrelated labels to compare the weight distribution for different layer sizes *M* and different correlation parameters *s* (**Supplementary Fig. 5**). For these models, the weights were found by solving the convex optimization problem with the *cvxopt* Python package.

#### Pattern classification

We define a classification task with *P* patterns as 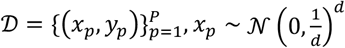, *y*_*p*_ ∈ {0,1}, where *y*_*p*_ is drawn *i*.*i*.*d*. The network is trained to predict the label *y*_*p*_ for any pattern *x*_*p*_. Predictors *W* are either initialized as a null matrix or each weight is drawn *i*.*i*.*d*. from 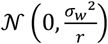, where *r* is the predictor’s parameter count (in-degree). Unless stated otherwise, 20 epochs of training are performed, i.e., each pattern is visited 20 times. The sequence of patterns is shuffled randomly for every epoch. In some cases, classifiers are trained on “noisy” versions of the patterns defined as 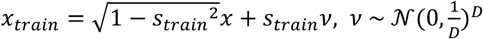. The parameter *s*_*train*_ determines the balance of signal and training noise in that for *s*_*train*_ = 1, the model is trained solely on noise.

During training, for each pattern, the network predicts a label. The prediction by the *i*-th ensemble member is given by *ReLU (f*_*i*_*(x*_*p*_)) = *ReLU (W*_*i*_ *diag(R*_*i*_*) ψ(x*_*p*_)). The ensemble prediction *f*_*ens*_*(x*_*p*_) is decided by majority vote (mode). We define a signed prediction error 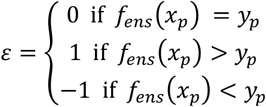. After each prediction, *W* is adapted according to the learning rule Δ*W*_*ij*_ = −*η ψ*_*j*_*(ϵ* + *β(f*_*i*_ − *f*_*ens*_).

The generalization performance of the network is evaluated by testing its prediction accuracy on “noisy” versions of the training patterns 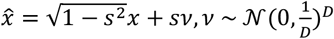 with generalization difficulty *s*. The error rate (false predictions divided by total predictions) is used as performance metric.

#### Regression

A regression task with *P* training patterns is given by 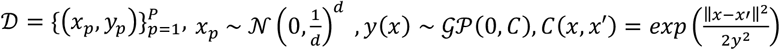. In brief, we uniformly sample inputs from the unit sphere and draw the corresponding targets from a Gaussian process, the covariance of which is defined by a Gaussian (RBF) kernel with length scale *γ*. In doing so, we generate a continuous, nonlinear mapping ℝ^*d*^ → ℝ. For further details on this task framework, see Xie et al. (2023). In some cases, we corrupt the training targets with Gaussian noise, so that *ŷ(x)* = *y(x)* + *ν, ν* ∼ *N(*0, *σ*^2^)^*D*^. For Ridge regression, we solve the Ridge minimization problem with the *scikit learn* Python package with intercept 0.

In the case of online learning, training pairs of inputs and outputs are presented to the predictors repeatedly. In each epoch, the weights are fit by Δ*W*_*ij*_ = −*η ψ*_*j*_*(x)(ε* + *β(f*_*i*_*(x)* − *f*_*ens*_*(x)))*, where *ε* = *f*_*ens*_*(x)* − *y(x). η* corresponds to the learning rate which is decreased during training if, for a given *η*, the mean training error over all patterns did not decrease thrice. Once *η* < 10^−6^, training is stopped. The weight modification term can be rearranged as Δ*W*_*ij*_ = −*η ψ*_*j*_*(x)(β f*_*i*_*(x)* + (1 − *β) f*_*ens*_*(x)* − *y(x))*. Here it becomes clear that for *β* = 1, all ensemble members are trained based on their individual error.

In **Fig. 4d**_**3**_, we computed the effective Ridge penalty of subsampling ensembles. To do so, we minimize the Euclidean distance between the output vectors of the ensemble and a fully connected Ridge predictor. The Ridge penalty of the minimizing predictor is then taken as the effective Ridge penalty. The optimal Ridge penalty is numerically found.

Since we are interested in weight regularization, we consider regression tasks in the overparameterized regime where *P* > *M*.

#### Centered kernel alignment

To measure the similarity between two subsets of the expansion layer (**Supplementary Fig. 4g**), we applied centered kernel alignment (CKA). In simple terms, to compare two feature spaces, CKA measures whether inputs that have a similar representation in the first feature space also have a similar representation in the second feature space. Precisely, CKA of two Kernels *K, K*’ is defined as their normalized Hilbert-Schmidt Independence Criterion (HSIC) (Gretton et al., 2005; Kornblith et al., 2019): 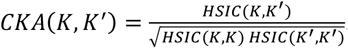. To calculate CKA, we computed the empirical linear Gram matrices of expansion layer subsets (*M* = 10000) over 15000 patterns of dimension *d* drawn uniformly from the unit sphere.

#### Statistical testing

To evaluate the null hypothesis that empirical connectivity follows a Bernoulli model, we used Monte Carlo tests. For a given statistic *T* with empirical value *T*_*empirical*_, we conduct a Monte Carlo test by computing the fraction *p* in *n* independent initializations of the Bernoulli model for which *T*_*Bernoulli*_ differs at least as much from its expected value under the null hypothesis *T*_*empirical*_.

To compare connectivity obtained from pruning models to empirical connectivity, we conducted permutation tests. For a given statistic *T*, the null hypothesis is that both empirical and modeled connectivity are derived from the same distribution. We exchange data points between empirical and model data and compute the fraction *p* in *n* shuffles for which *T* of the shuffled distributions is at least as large as *T* of the two original distributions.

## ACKNOWLEDGEMENTS

We thank Nate Sawtell, Ashok Litwin-Kumar, Nicolas Brunel, and Wade Regehr for comments on a previous version of this manuscript. A.H. is grateful for funding from Boehringer Ingelheim Fonds. R.K. acknowledges support from the William Randolph Hearst Fund. This work was supported by NIH grants RF1MH128949 and RF1MH129261 to W.C.A.L.

