## Supplementary figures for "Random innervation of cerebellar Purkinje cells as a substrate for diverse representational learning"

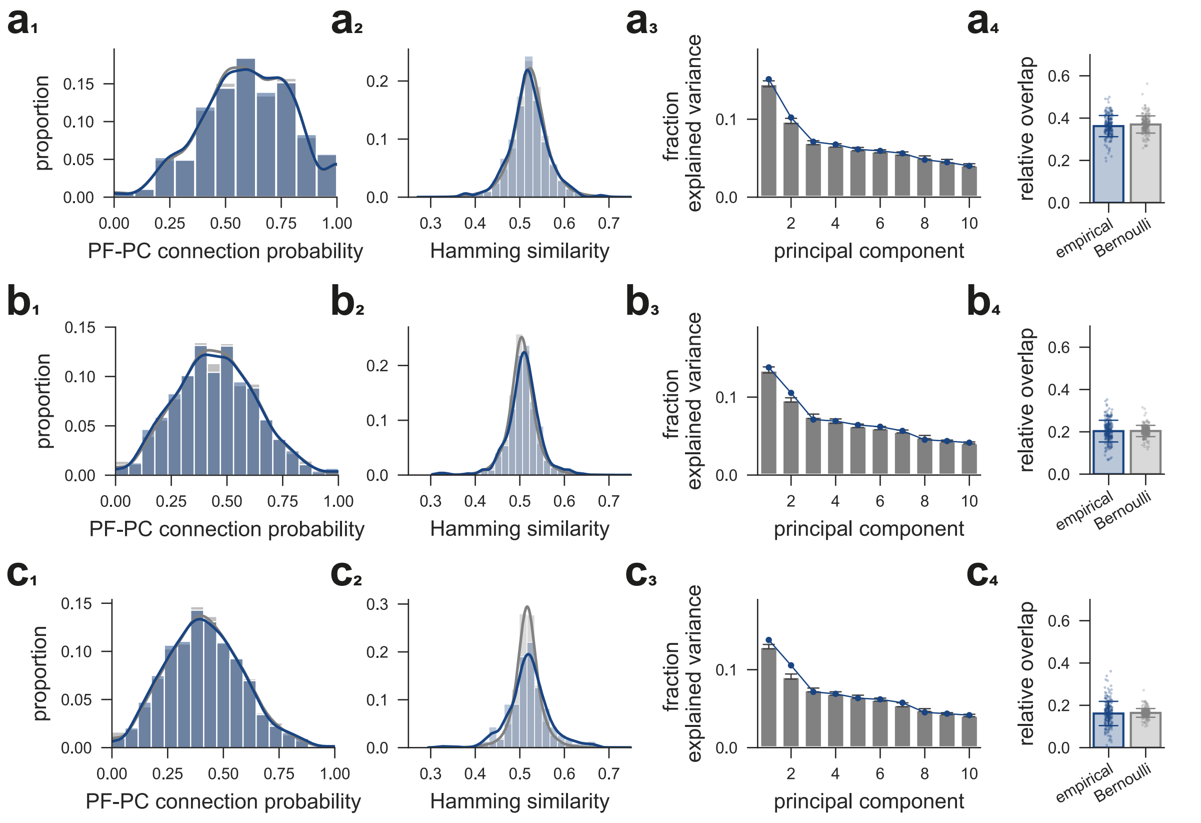


**Supplementary Fig. 1**

**a)** Connectivity metrics similar to those in **Fig. 2** for a touch threshold of 16nm. **b)** Same as in **a** except for a touch threshold of 500nm. **c)** Same as in **a** except for a touch threshold of 10000nm. Note how the variance in Hamming similarity and in relative overlap becomes inflated. This is expected if PFs are considered that are trivially unconnected due to their spatial displacement.

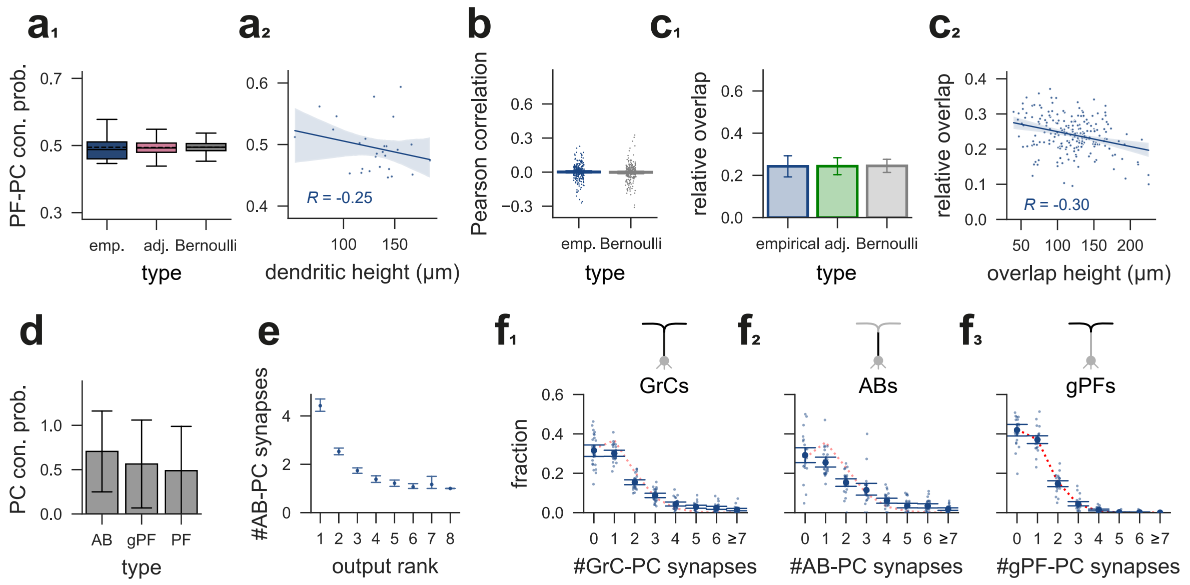


**Supplementary Fig. 2**

**a$_1$)** PF-PC connection probability for PCs. The variance of the distribution is slightly elevated compared to the Bernoulli model, indicating that some PCs may systematically sample fewer PFs. This is partially due to the positioning of their dendritic tree in the molecular layer (**a$_2$**). An adjusted model corrected for this effect is shown in the middle (adj.). Dashed lines indicate the average. **a$_2$)** PF-PC connection probability for PCs as a function of the average height of their PF touches. Since PF-PC connection probability shows a weak dependence on height in the molecular layer (Nguyen et al. 2023), PCs with more dendritic branches at the top are expected to have a slightly reduced connection probability. This explains some of the elevated variance observed in **a$_1$**. **b)** Pairwise Pearson correlation of PF input vectors on shared touches for pairs of PCs. Only PC pairs with at least 30 shared touches were included (error bars: mean ± 95% CI). **c$_1$)** PF relative input overlap (shared PFs/shared touches) for pairs in **b**. Note that the variance of the empirical connectivity is slightly larger compared to the Bernoulli model. An adjusted Bernoulli model that retains the connection probability of each PC is shown in green. This model demonstrates that the elevated variance of the empirical data is largely explained by the increased variance of PF-PC connection probabilities in **a$_1$** (see also **Supplementary Fig. 1**, error bars: mean ± 95% CI). **c$_2$)** Relative PF input overlap for pairs in **c$_1$** as a function of the average height of their shared PF touches (overlap height, see **a$_2$**). If two PCs share most touches high in the molecular layer, they are expected to have a lower connection probability to these shared touches and hence a lower relative overlap. This explains some of the increased variance in **a$_2$**. **d)** Isolated PC connection probability for ABs and gPFs in cases where only the AB (the gPF) but not the gPF (AB) of the same GrC touch a given PC. The connection probability of remote PFs is shown for comparison (error bars: mean ± SD). **e)** AB-PC synapse count for the top-n outputs of each AB. **f$_1$)** Analogous to **Fig. 1e** but for local GrCs. The optimal fit of the Poisson-Bernoulli combined model does not reproduce the synapse count distribution (error bars: mean ± 95% C.I). **f$_2$)** Analogous to **f$_1$** but for ABs. **f$_3$)** Analogous to **f$_2$** but for gPFs. Here, as for remote PFs, the Poisson-Bernoulli combined model does reproduce the synapse distribution.

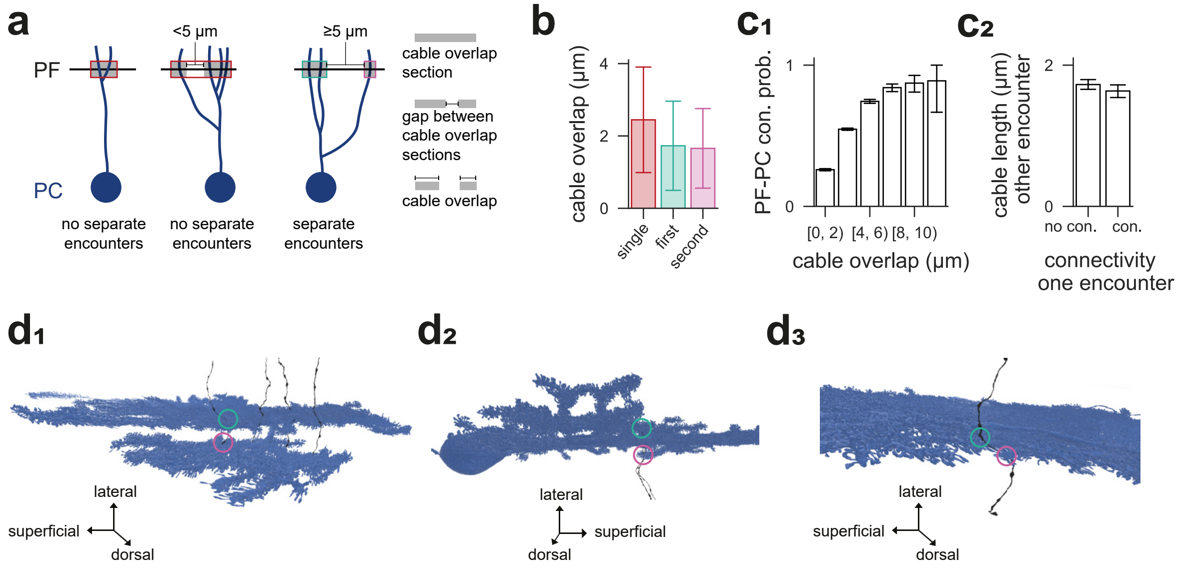


**Supplementary Fig. 3**

**a)** Schematic of individual PF-PC pairs, considered to have no separate encounters (left) and two separate encounters (right). An encounter is defined as a location where the PF and the PC dendrites are in a spatial proximity of 160 nm (cable overlap section; gray shaded areas). Two such encounters are considered separate, i.e., spatially independent, when the distance between them exceeds a threshold of 5 µm along the transversal axis (Methods). The cumulative length of these sections (along the transversal axis) is a proxy for the intensity of a PF-PC touch (cable overlap). **b)** Cable overlap, as defined in **a,** for touching PF-PC pairs with no separate encounters (single) and two separate encounters (first, second). Each of the two separate encounters in the latter case typically show smaller cable overlap than the single encounters in the former case (error bars: mean ± SD). **$c_1$)** The PF-PC connection probability is strongly correlated with the cable overlap of the PF-PC touch. Together with **b**, this explains the average connection probability of ~1/3 for each individual of the two separate encounters observed in **Fig. $2f_3$**. (error bars: mean ± 95% CI). **$c_2$)** Correlations between a connection at one encounter and the cable overlap of the second encounter are negligible. Hence the analysis in **Fig. $2f_3$** is not confounded by an influence of connections at one encounter on cable overlap at the other encounter ($R$ = -0.04, error bars: mean ± 95% CI). **d)** Example reconstructions of PF-PC pairs with separate encounters (green and magenta).

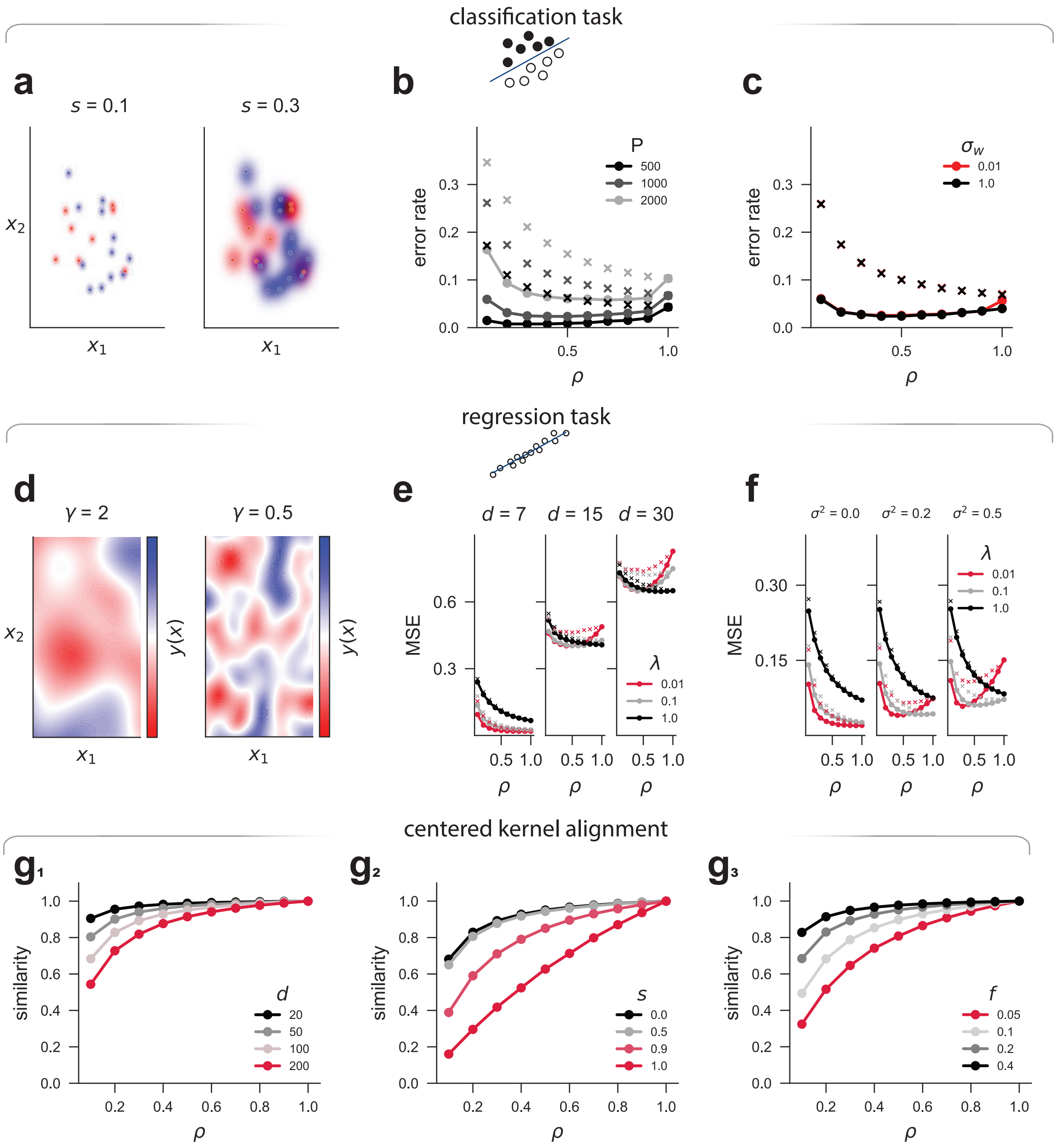


**Supplementary Fig. 4**

**a)** Classification task: Patterns are sampled from a Gaussian distribution and are assigned a random label (blue or red). A network learns to separate blue from red patterns (dots). Generalization performance is then examined using corrupted versions of the patterns, where the signal is partially replaced by Gaussian noise (contours). The signal-to-noise ratio is controlled by parameter $s$ (Methods). **b)** Generalization error rate in a classification task with uniform weights for different numbers of patterns $P$. The setup is the same as in **Fig. 5a₁**. **c)** Generalization error rate with initial Gaussian weight distribution. $\sigma w$ controls the spread of the Gaussian (Methods). Note that diversity in initial (unstructured) weights reduces the benefit of subsampling. **d)** Regression task: Inputs are sampled from a Gaussian distribution, and target values are generated by a Gaussian process the covariance of which is defined by a radial basis function kernel (Methods). The length scale $\gamma$ controls the smoothness of the target function. The network learns to approximate the target function based on randomly drawn examples and is then tested on unseen pairs of input and target output. **e)** Extension of **Fig. 5d₂**: Mean-squared generalization error in Ridge regression as a function of $\rho$. Top: different task dimensions $d$. Bottom: different noise levels $\sigma$ in the training targets. $\lambda$ denotes the explicit regularization penalty. Subsampling ensembles (partial connectivity) display implicit regularization that adapts to the task. With low explicit regularization (red), subsampling ensembles show stronger performance for both low- and high-dimensional tasks (see red arrowheads). This advantage is also evident across different noise levels. **f)** Same as **e** but for different target noise $\sigma$. Tasks with high target noise require stronger generalization. **$g_1$)** Centered kernel alignment (CKA) of two expansion layer subsets for different degrees of partial connectivity and task dimensions. CKA measures the similarity of two feature representations (PF subsets in this case) that serve as the input to two Purkinje cells. For $\rho = 1$, PF inputs to PCs are identical and the similarity is 1. In the case of partial connectivity, similarity originates from input overlap as well as correlations across PFs. The latter depends on the input structure. High input dimension makes PFs more dissimilar. This information is retained by partially connected ensembles and explains their adaptive regularization behavior (**f**). **$g_2$)** Same as **$g_1$** but for a signal to noise parameter $s$ in the expansion layer. For $s = 0$, the expansion layer is unperturbed. For $s = 1$, the expansion layer encodes Gaussian noise. **$g_3$)** Same as **$g_1$** but for the fraction of active neurons $f$ in the expansion layer. Sparsity makes the subsets more diverse.

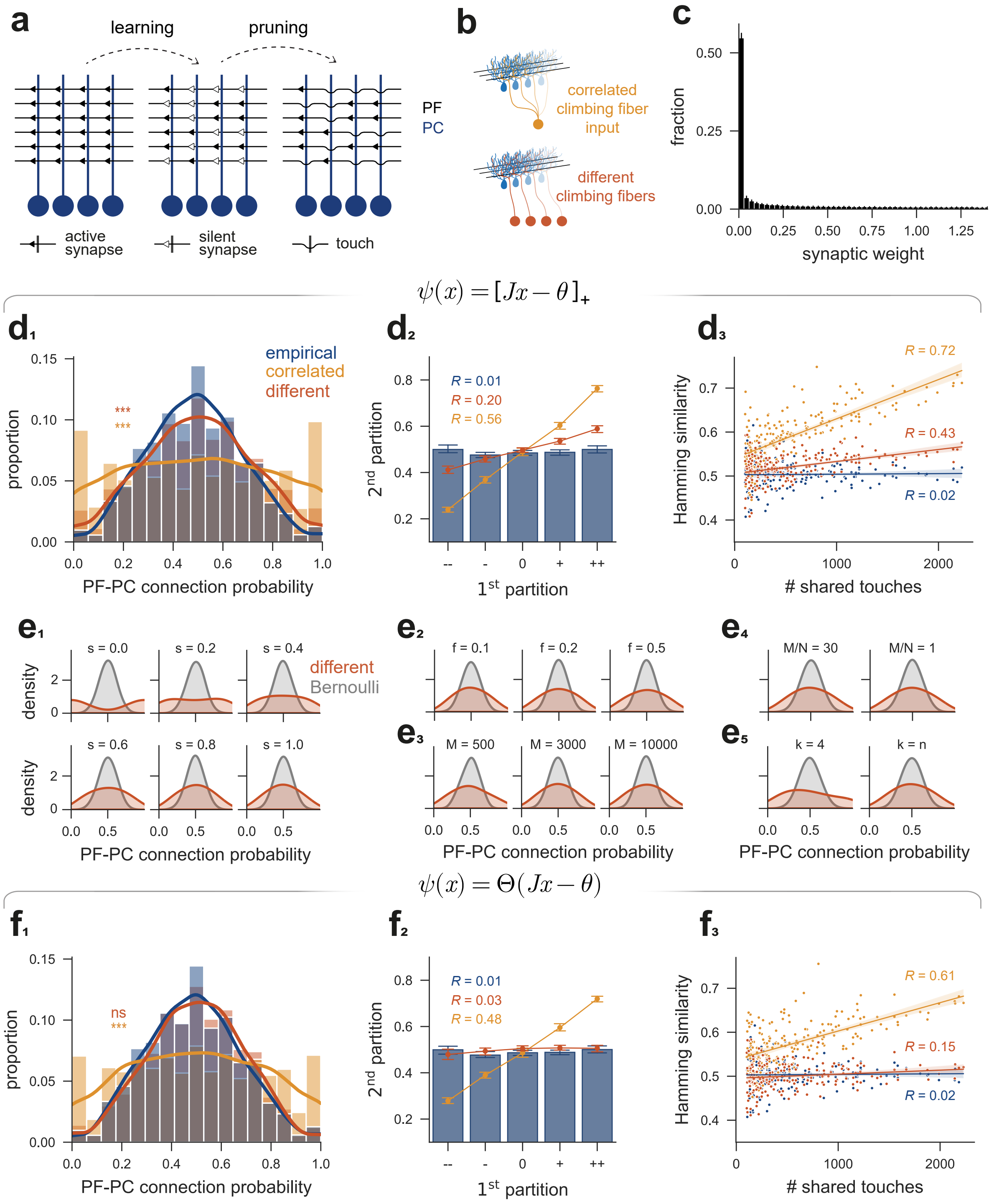


**Supplementary Fig. 5**

**a)** Schematic model of learning-induced synaptic pruning. Each PC learns to associate PF input patterns with binary labels. At maximum storage capacity, a considerable fraction of PF-PC synapses becomes silent. In an alternative explanation of partial connectivity, pruning of silent synapses would result in PFs passing PC dendrites without synapsing, that is, what we observe as touches. **b)** Schematics of two extreme cases where labels are the same for all PCs (shared climbing fiber; orange) or fully uncorrelated between PCs (different climbing fibers; red). In either case, PCs in our model share PF input as determined by their shared touches (Methods). The color convention for conditions is maintained throughout. **c)** Synaptic weight distribution obtained from storing information in strictly non-negative PF-PC synapses (Brunel et al. 2004). Note that about 50% of the weights vanish, i.e., the synapses are silent. **$d_1$)** PF-PC connection probabilities as in **Fig. 2$d_1$** for the true connectivity and the two variants of the pruning model. If the PF code is obtained by rectified-linear thresholding, in both the correlated and the uncorrelated scenarios, the resulting connectivity deviates significantly from the empirical one (Permutation test for difference in variance, $p < 0.001$). **$d_2$)** Cross validation of PF connection probabilities by random bipartitioning of postsynaptic PCs. For each PF, the touched PCs were randomly divided into two balanced partitions. PFs were then sorted into 5 bins depending on their connection probability to the first partition (minimal: --, maximal ++). Connection probabilities to the second partition are shown on the y-axis. Note that for the pruning models, but not for the true connectivity, connection probabilities to the first subset are predictive for those to the second subset (empirical connectivity: Pearson $R = 0.01$, uncorrelated: $R = 0.20$, correlated: R = 0.56, error bars: mean ± 95% CI). **$d_3$)** Hamming similarity of pairs of PCs as a function of their shared touches. For the pruning models, overlap in touches (i.e., overlap in input space) results in larger agreement in sampling decisions. This is not the case for the empirical connectivity. Included are pairs of PCs with at least 100 shared touches. **$e_1$)** Version of the pruning model where 20 artificial PCs are connected to 3000 PFs. PCs learn associations with different degrees of label correlations ($s = 0$: identical, $s = 1$: fully uncorrelated). Silent synapses are pruned afterwards. **$e_{2\text{-}4}$)** Same setup as in **$e_1$** but across different parameters at $s = 1$: PF coding sparsity $f$ (**$e_2$**), PF layer size $M$ (**$e_3$**), expansion ratio $M/N$ (**$e_4$**), and PF in-degree $k$ (**$e_5$**). The finding that certain PFs are over- or under-represented, despite no correlations between storage labels, is robust. **$f_{1\text{-}3}$)** Same as **$d_{1\text{-}3}$**, but for a model where the PF code is fully binary. Here, in the uncorrelated case, most of the structure is lost (different vs. empirical: Permutation test, $p = 0.30$; correlated vs. empirical: Permutation test, $p < 0.001$).
